# Developmental enrichment reorganizes the directional topography of mouse proprioceptive cortex

**DOI:** 10.64898/2026.07.14.738599

**Authors:** Melanie Palacio-Manzano, Fei Ma, Coralie Hoffmann-Schreiner, Yann Ravussin, Mario Prsa

## Abstract

The organization of proprioceptive cortex emerges from experience-dependent patterns of limb use during development, rather than merely reflecting peripheral innervation. However, the degree and specifics of these developmental changes remain poorly understood. To address this question, we investigated how early environmental enrichment (EE) reshapes the topography and functional properties of the mouse forelimb proprioceptive cortex. Using wide-field calcium imaging, we found that EE did not induce major reorganization of the mesoscale activation map. To resolve cellular level topography, we used a custom large-field fluorescence macroscope enabling unbiased single neuron imaging across the entire activation area and mapped the preferred limb movement direction of each neuron. We identified a topographically organized directional bias (anisotropy) in neuronal tuning for movements directed toward the body, with neurons in somatosensory and motor parts of the map preferring different movement directions. EE did not abolish this anisotropy but reorganized it. Anisotropic zones matched the somatosensory and motor anatomical locations more closely, and neurons in both areas aligned to a single preferred direction. Our findings therefore show that enriched limb use, rather than redrawing the cortical map, coordinates its sensory and motor components, revealing a developmental mechanism that tunes the mouse proprioceptive cortex for body-directed movements prominent in its natural behavior.

## Introduction

The use of hands and limbs is central to our existence, driving everything from basic object manipulation to complex social interactions. The sensory aspects of this behavior comprise two distinct modalities: tactile feedback for processing external touch stimuli, and proprioception for tracking own movement and posture. The primate primary somatosensory cortex processes these inputs separately, mapping tactile signals to area 3b and proprioceptive signals to the more rostral area 3a, with a similar architectonic distinction found across other mammalian brains[1–9].

These two areas differ fundamentally in their topography and development. Tactile area 3b features a precise topographic map with small receptive fields, where densely innervated limb structures, like the digits, occupy larger cortical territories[2, 6]. In fact, it often seems to form an isomorphic three-dimensional representation of the periphery[10–14] and appears organized and functional at birth[15]. In contrast, proprioceptive area 3a remains largely non-functional at birth[15] and exhibits distinct configurations across species that use their limbs and hands differently[6]. Rather than being hard-wired, the proprioceptive cortex appears to emerge during development to reflect behaviorally relevant specializations, varying even among individuals of the same species[6].

In the mouse forelimb somatosensory cortex (fS1), we previously found that tactile and proprioceptive maps are not strictly segregated[16]. Moreover, individual proprioceptive neurons exhibit a striking functional anisotropy, overrepresenting limb displacements toward the body, into the peripersonal space, while underrepresenting body-fugal movements[16]. Given that the proprioceptive cortex emerges postnatally to reflect behavioral repertoires, we hypothesize that these peculiar anatomical and functional features may result from development within the restrictive, standard housing conditions of laboratory mice. Having been bred for experimental use and effectively isolated from feral conditions for over 100 years (since the 1920s), the C57BL/6 genetic strain may exhibit cortical wiring adapted to this lack of sensorimotor and environmental diversity.

To test this hypothesis, we reversed environmental impoverishment by introducing enriched sensorimotor experiences during development. Since Donald Hebb’s pioneering 1940s studies on the role of early experience on learning[17], such environmental enrichment has been known to exert widespread effects on the brain, from macro-scale morphological changes to cell-level structural and functional plasticity[18]. Accordingly, here we studied how environmental enrichment shapes the organization of the fS1 proprioceptive map and the tuning properties and topography of its neurons.

We find that directional anisotropy is a specialization of the mouse proprioceptive cortex that is topographically malleable across developmental conditions. It adapts to environmental complexity within a globally stable cortical architecture optimized for peripersonal movement feedback.

## Results

### Environmental enrichment induces metabolic and behavioral changes

To assess eventual reorganization of the proprioceptive cortex during development, we compared mice raised in conventional housing (CH) and an enriched environment (EE). The EE mice were reared in a specialized habitat designed to promote increased locomotor activity, climbing and skilled forelimb engagement through reach-to-grasp training and fine manipulation (Video 1, see Methods). Because EE paradigms are inherently multifactorial and non-standardized[18], we first verified that our specific protocol produced expected changes in mouse physiology and physical phenotype.

At 15 weeks of age, automated metabolic and behavioral monitoring as well as body composition analysis revealed phenotypic differences between the two groups. EE mice exhibited significantly lower body weights and a lower percent body fat compared to CH controls, alongside a reduced metabolic rate (Fig. 1A, B). To determine whether this metabolic difference was driven by differences in body composition, we performed an ANCOVA with fat mass and fat-free mass as covariates. During the diurnal period, the group effect was no longer significant after controlling for lean and fat mass (p = 0.45), indicating that daytime metabolic differences are fully accounted for by body composition. During the nocturnal period, however, the group effect remained significant (p = 0.018) and was accompanied by higher food and water consumption and increased locomotor activity in EE mice (Fig. 1C, D). This indicates that EE mice achieve lower nocturnal energy expenditure despite being more physically active. Because this effect persists even after accounting for body composition, it points toward a significant improvement in metabolic efficiency.

**Figure 1.**
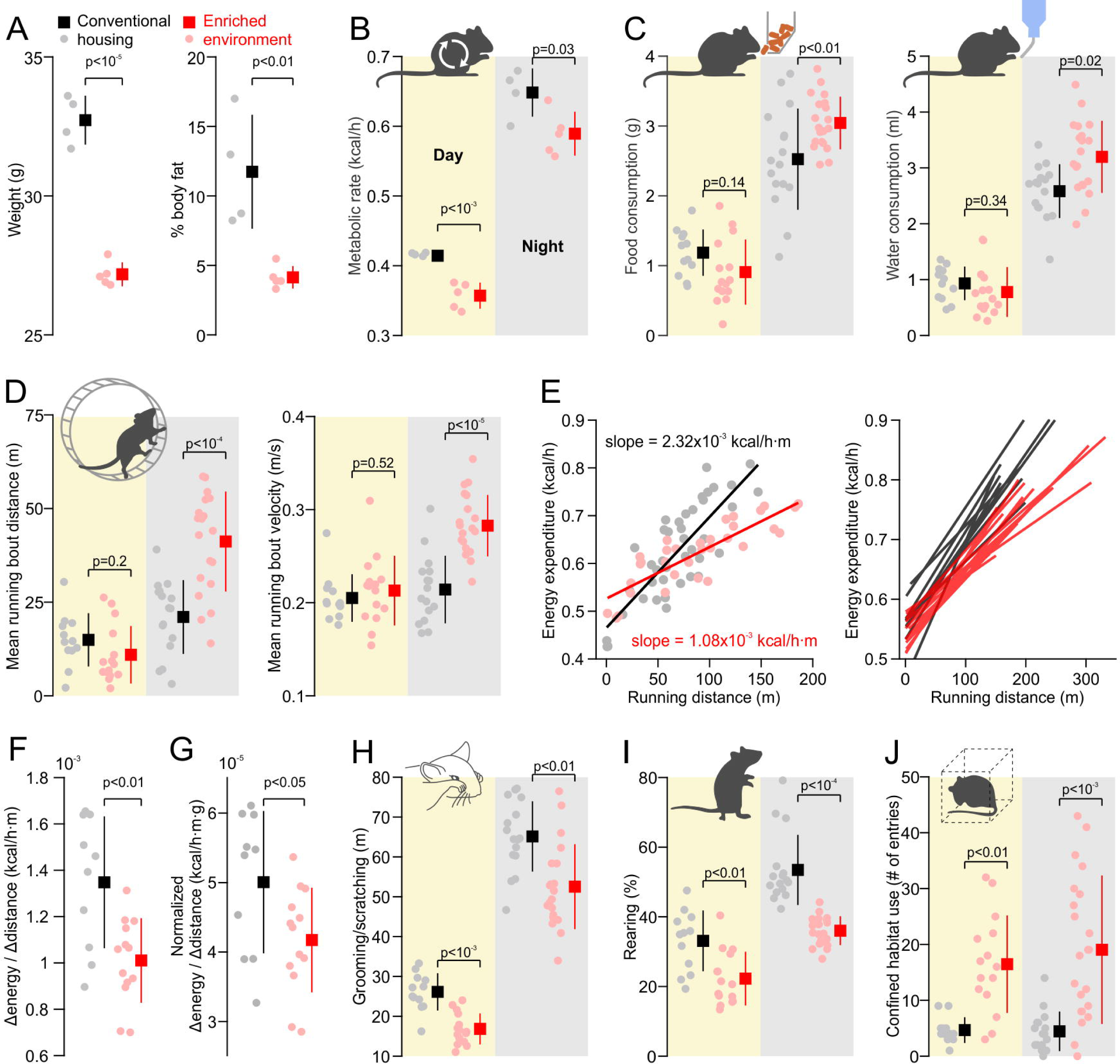
Metabolic and behavioral effects of environmental enrichment. **(A)** Weight and body composition (% fat mass) comparison between EE (N=5) and CH (N=4) mice (two-sample *t* test). Shaded dots are data of individual mice. Squares and delimiters are means +/- s.d. **(B)** Metabolic rate comparison during diurnal (yellow shade) and nocturnal (grey shade) periods (data as in A). **(C)** Food and water consumption comparison (linear mixed effects model). Shaded dots are individual diurnal (yellow shade, N=12 sessions for 4 CH mice and N=15 sessions for 5 EE mice) and nocturnal (grey shade, N=16 sessions for CH and N=20 sessions for EE) sessions across mice. Squares and delimiters are means +/- s.d. **(D)** Comparison of wheel running activity (data as in C). **(E)** Left panel: Energy expenditure as a function of running distance on the wheel for an example 24h session of a CH (black) and EE (red) mouse with linear regression fits. Measurements (shaded data points) correspond to individual running bouts. Right panel: linear regression fits for all individual 24h sessions (nocturnal and diurnal data pooled; 10 sessions for CH and 15 sessions for EE) show consistently lower slopes for EE mice. **(F)** Comparison of metabolic efficiency slopes in E (linear mixed effects model). **(G)** Same data and comparison as in F after normalizing for individual fat free mass. **(H, I, J)** Comparison of the amount of detected fine movement (grooming and scratching), % of time spend in the rearing position (outside of wheel use, drinking, feeding and ambulatory periods) and number of entries into a confined habitat (linear mixed effects model). All data as in C.

To quantify differences in metabolic efficiency, we analyzed the relationship between energy expenditure and physical output during active periods containing wheel-running bouts (Fig. 1E). EE mice showed consistently lower slopes (Fig. 1E,F) meaning that increases in their motor activity had a lower energetic cost compared to CH mice. Notably, the increased efficiency persisted after normalizing for individual fat-free mass (Fig. 1G). This physiological shift suggests an adaptation in muscle fiber composition, specifically an enrichment of Type I (oxidative) fibers which optimize ATP production per unit of oxygen.

Furthermore, EE reduced excessive behavioral stereotypies such as grooming/scratching (Fig. 1H) and prolonged rearing (Fig. 1I), while promoting exploratory behaviors, as evidenced by increased entries into a confined habitat (Fig. 1J). These differences likely reflect EE’s well-documented anxiety-reducing effects[19–22].

Because it has been reported that EE induces structural and functional changes in visual [23] and somatosensory[24] cortices, and that exercise specifically modifies propriospinal neuron subtype composition[25], we next tested whether the observed physiological and behavioral shifts translate into a reorganization of the proprioceptive cortex.

### Environmental enrichment fails to reorganize the mesoscale cortical proprioceptive map

To stimulate proprioceptive afferents, we passively displaced the mouse forelimb with a robotic manipulandum following our established protocol[16]. Using wide-field calcium imaging, we simultaneously imaged cortical activation in transgenic mice expressing the calcium indicator GCaMP6f in L2/3 neurons (Fig. 2A). Consistent with our previous results[16], the proprioceptive stimulation activated a cortical area spanning the primary forelimb somatosensory (fS1) and motor (CFA: caudal forelimb area) cortices in the CH (N=9 mice, 45 sessions) and EE (N=7, 38 sessions) mouse groups (Fig. 2B).

**Figure 2.**
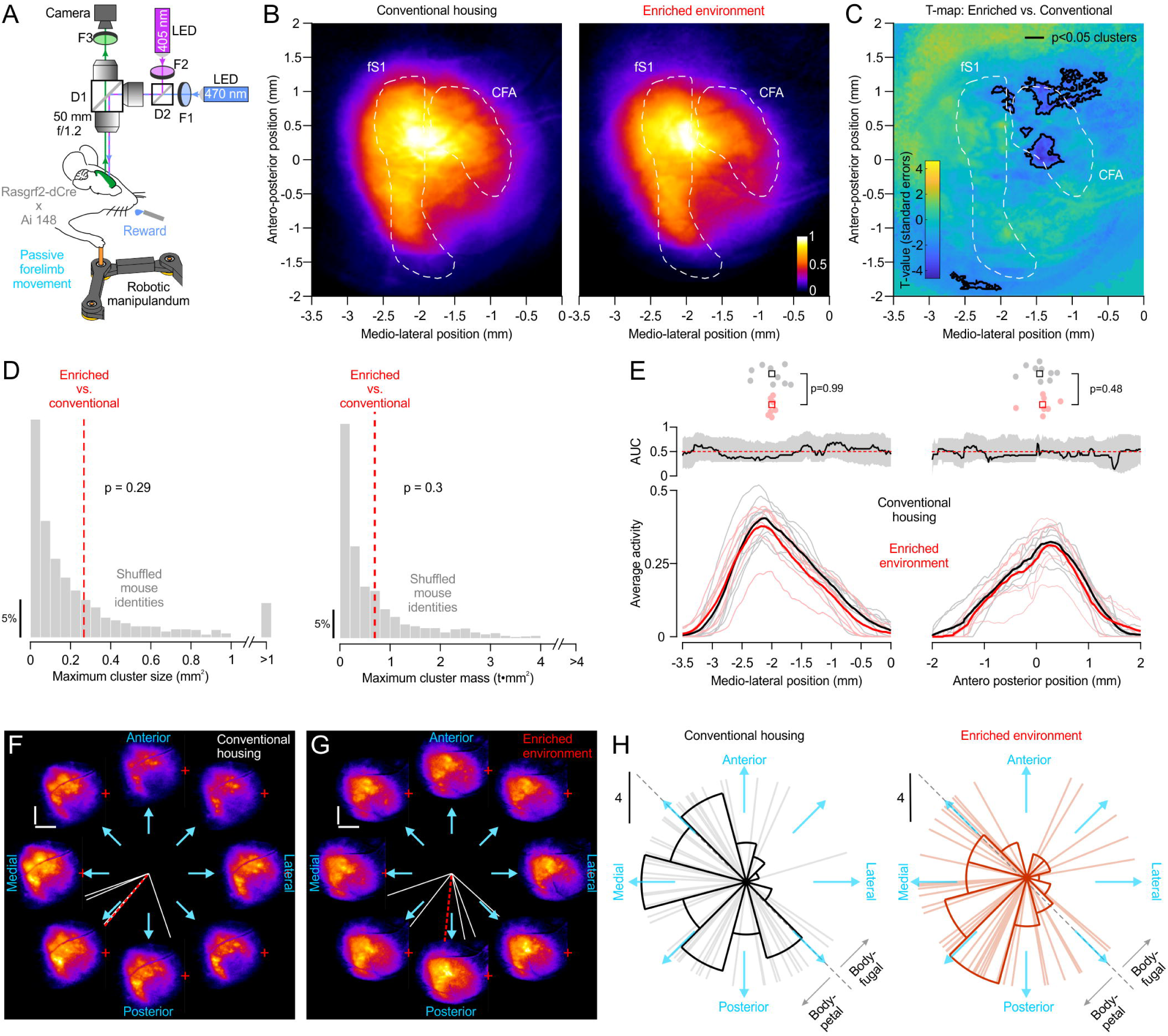
Topographic and functional stability of the cortical proprioceptive map. **(A)** Wide-field population imaging of L2/3 GCaMP6f activity during forelimb proprioceptive stimulation (D1, D2: dichroic mirrors; F1, F2, F3: bandpass filters). **(B)** Average population GCaMP6f cortical activation by contralateral forelimb proprioceptive stimulation (normalized to peak activity) for CH (N=9, 45 sessions) and EE (N=7, 38 sessions) mice. The (0,0) origin is the location of bregma. The fS1 and CFA contours are based on locations of cortico-spinal projection neurons retrogradely labeled from the cervical cord and forelimb muscles[38]. **(C)** T-map comparison of proprioceptive activation maps across CH and EE mice. **(D)** Comparison of maximum size and mass of significant t-map clusters, between the empirical differences (red dotted lines) and shuffled null distributions. P-values (permutation test) correspond to the fraction of null distribution values more extreme than the empirical data values. **(E)** Bottom: average activity along the medio-lateral and antero-posterior dimensions of the cortical surface (shaded lines: individual mice; bold lines: population means). Middle: AUC values for each sampled location from the ROC analysis (black trace). The 95% confidence intervals (shaded areas) were not more extreme than the chance 0.5 discrimination level (dotted red lines). Top: Comparison of peak activity locations (two-sample *t* test; shaded dots: individual mice; square: population means). **(F, G)** Activation maps for each stimulus direction (cyan arrows) in representative CH and EE mouse sessions. White lines denote the preferred direction of individual sessions, while the red dashed line represents the resultant preferred direction averaged across all sessions for that mouse. **(H)** Distribution of population activity preferred directions across all imaging sessions (individual lines; N=10 CH mice, 51 sessions; N=8 EE mice, 43 sessions).

To evaluate topographic differences in the cortical proprioceptive maps between the two groups, we generated a t-statistic map representing the signal-to-noise ratio of the group difference at each pixel (see Methods). We identified difference clusters (based on a t-statistic threshold corresponding to p<0.05) primarily in the CFA region (Fig. 2C). To determine if these clusters occurred by chance, we performed a permutation test by randomly shuffling the group labels to construct null distributions for maximum cluster size and maximum cluster mass (Fig. 2D). These two measures for the detected clusters in Fig. 2C exceeded only ∼70% of the null-distribution values, meaning that we cannot statistically rule out that they occurred by chance.

To compare the spatial features of the proprioceptive activation maps more directly, we next ran a Receiver Operating Characteristics (ROC) analysis on the average activity samples along the medio-lateral and antero-posterior dimensions of the cortex. The area under the curve (AUC) measure did not exceed the 95% bootstrapped confidence intervals at any location (Fig. 2E). The groups also showed no difference in the size of the total activation area (10.6 ± 1.4 mm^2^ for CH vs. 9.2 ± 2.3 mm^2^ for EE, p=0.17, non-paired t-test) or in their peak activity locations (Fig. 2E, top).

These findings suggest that environmental enrichment does not drive major topographic reorganization of the proprioceptive cortex at the mesoscale level, pointing instead toward potential functional alterations. Therefore, we next assessed the directional tuning of the population activity. Consistent with our previous single-cell observations[16], the macro-level population response was directionally tuned, exhibiting an overt preference for limb displacements toward the peripersonal space (i.e. in medial and posterior directions). Although we hypothesized this anisotropy might diminish in EE mice due to their increased exploratory limb use in larger spaces, the distribution of preferred directions across sessions remained non-uniform in both groups (Fig. 2H, p<0.001, Rayleigh test). Actually, the non-uniformity appeared qualitatively enhanced in the EE group (Fig. 2H), though the concentration of the two circular distributions did not statistically differ (p=0.41, Wallraff test).

Since neither macro-level topography nor population tuning exhibited major changes following EE, we next sought to determine if functional differences instead manifest at the higher spatial resolution of single neuron activity.

### Proprioceptive anisotropy is topographically organized in the mouse cortex

To record single cell activity and obtain an unbiased sampling across the whole proprioceptive map we used a custom fluorescence macroscope[26] for imaging L2/3 neurons (Rasgrf2-dCre x Ai148 transgenic mice) in fields of view (∼9 mm^2^) covering entire fS1, CFA and surrounding cortical areas (Fig. 3A-B). A high-NA, low-magnification objective enabled the resolution of individual neurons using one-photon excitation of GCaMP6f (Fig. 3B-C). We identified 1162 neurons with proprioceptive responses (N=9 mice, 5 EE and 4 CH) by focusing the macroscope at different depths in L2/3 across imaging sessions. In agreement with the population activation map (Fig. 2B), proprioceptive neurons were predominantly located in fS1 (peak occurrence at ∼2.2 mm lateral to bregma) but also more medially in the CFA motor area (Fig. 3D).

**Figure 3.**
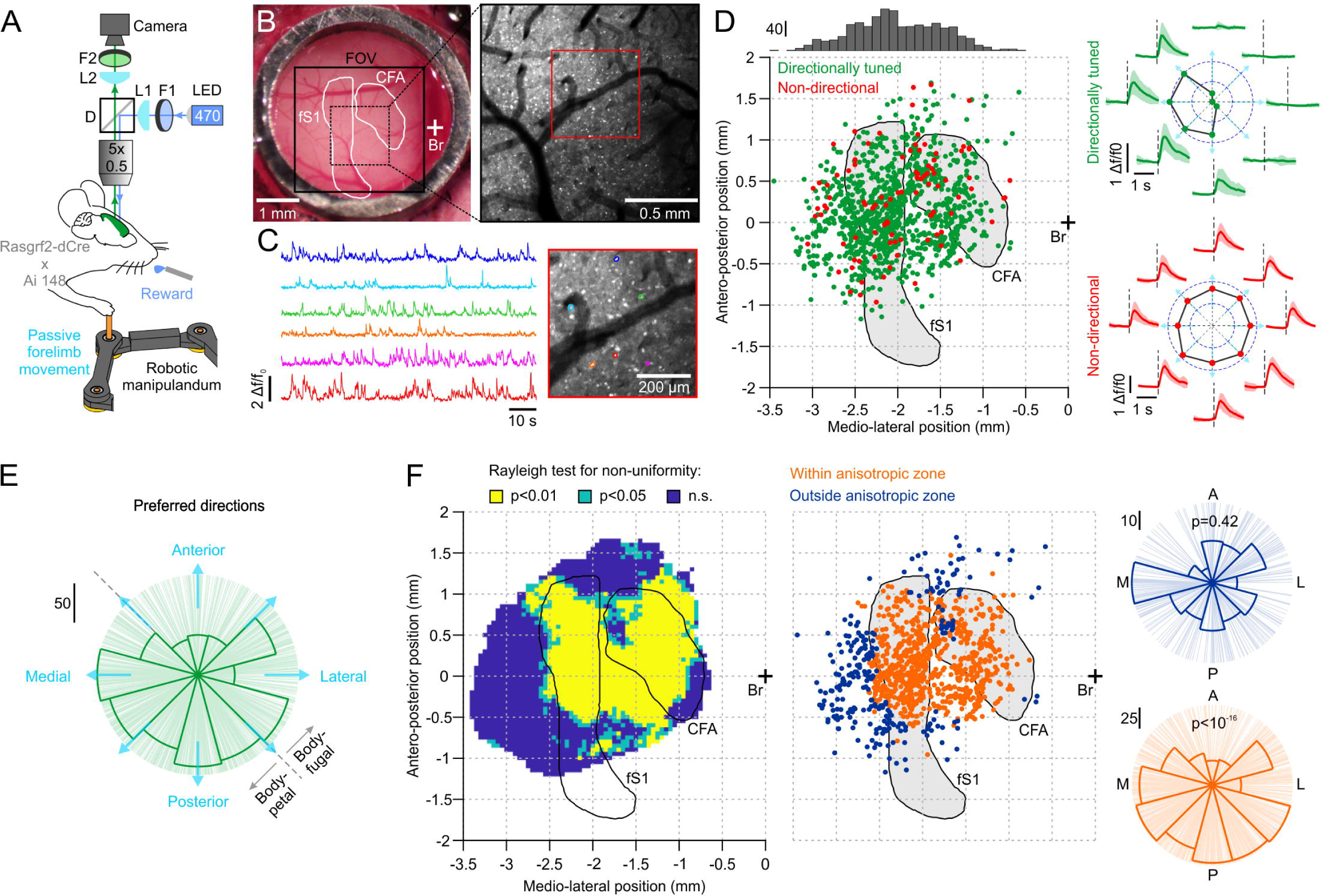
Topographic organization of the proprioceptive directional bias. **(A)** Wide-field single-cell imaging of L2/3 GCaMP6f activity during forelimb proprioceptive stimulation (D: dichroic mirrors; L1: focusing lens; L2: tube lens; F1, F2: bandpass filters). **(B)** Right: cranial window in the contralateral cortex of a representative mouse with fS1 and CFA contours (based on cortico-spinal tracing data [38]), bregma location (Br) and the imaging field of view (FOV). Left: maximum projection image of a FOV section showing the resolution of individual cell soma. **(C)** Example Δf/f_0_ activity traces (left) and corresponding spatial footprints (right) of detected individual neurons within the field of view highlighted by the red square in (B). **(D)** Left: spatial location of all imaged neurons with significant proprioceptive responses (N=1162, 9 mice) and their distribution along the medio-lateral axis of the cortex (top). Directionally tuned neurons (N=1067, significant von Mises fit) are depicted in green and non-directional neurons in red (N=95). Right: directional tuning curves for a representative directionally tuned (green) and a non-directional neuron (red). Green and red traces: mean (± s.d.) responses to the eight different movement directions (cyan arrows) Polar plots: peak activity as function of movement direction. Dotted lines: stimulus onset. **(E)** Distribution of preferred directions for all directionally tuned neurons (N=1067, individual lines). **(F)** Left: topographic representation of the directional anisotropy (p-values of the Rayleigh test for non-uniformity of preferred directions, see text for details). Middle: spatial locations of neurons in the uniform (blue, N=261) and non-uniform anisotropic (orange, N=806) directional tuning areas. Right: preferred direction distributions of the two groups with p-values (Rayleigh test).

By evaluating neural responses to limb displacements in the eight coplanar directions, we found that ≈92% (N=1067) of individual units were directionally tuned (Fig. 3D, in green), as determined by significant von Mises fits (see Methods). As observed previously using targeted two-photon imaging restricted to fS1[27], this unbiased population exhibited distinct tuning anisotropy (p<10^-14^, Rayleigh test) characterized by a single mode (p=0.21, excess-mass test for more than one mode) overrepresenting body-petal directions (Fig. 3E).

While these data establish a global preference for movements towards the peripersonal space, the critical advantage of mapping the entire proprioceptive cortex without sampling bias is the ability to resolve the spatial distribution of this preference. We therefore asked whether the anisotropy is homogenous across the entire map or if it exhibits a particular topographic organization. We spanned the 2D map with a circular sliding window (0.8 mm diameter, 0.05 mm steps) and performed a Rayleigh non-uniformity test for directionally tuned cells falling inside the window at each location. The resulting map of p-values (Fig. 3G) reveals that the statistically significant regions tend to be within the anatomical boundaries of fS1 and CFA, whereas the surrounding areas harbor neurons with uniformly distributed preferred directions.

Grouping the neurons located in the “significant” anisotropic and “non-significant” isotropic clusters confirmed the presence (p<10^-16^, Rayleigh test) and absence (p=0.42) of anisotropy within these two respective subpopulations (Fig. 3G, right), demonstrating that the directional bias is localized within a distinct cortical topography.

### Environmental enrichment modifies proprioceptive coding and its cortical topography

To determine whether environmental enrichment induces changes of cellular level activity and in its topographic organization, we analyzed the single-neuron tuning and their spatial distribution in EE (N=5) and CH (N=4) mice independently.

Neurons from EE mice were activated more reliably by the proprioceptive stimuli, exhibiting a significantly reduced coefficient of variation in their peak Δf/f_0_ responses (Fig. 4A). Concurrently, their directional selectivity decreased (Fig. 4A, lower κ in the von Mises fits), indicating that their responses were more broadly tuned across the eight movement directions.

**Figure 4.**
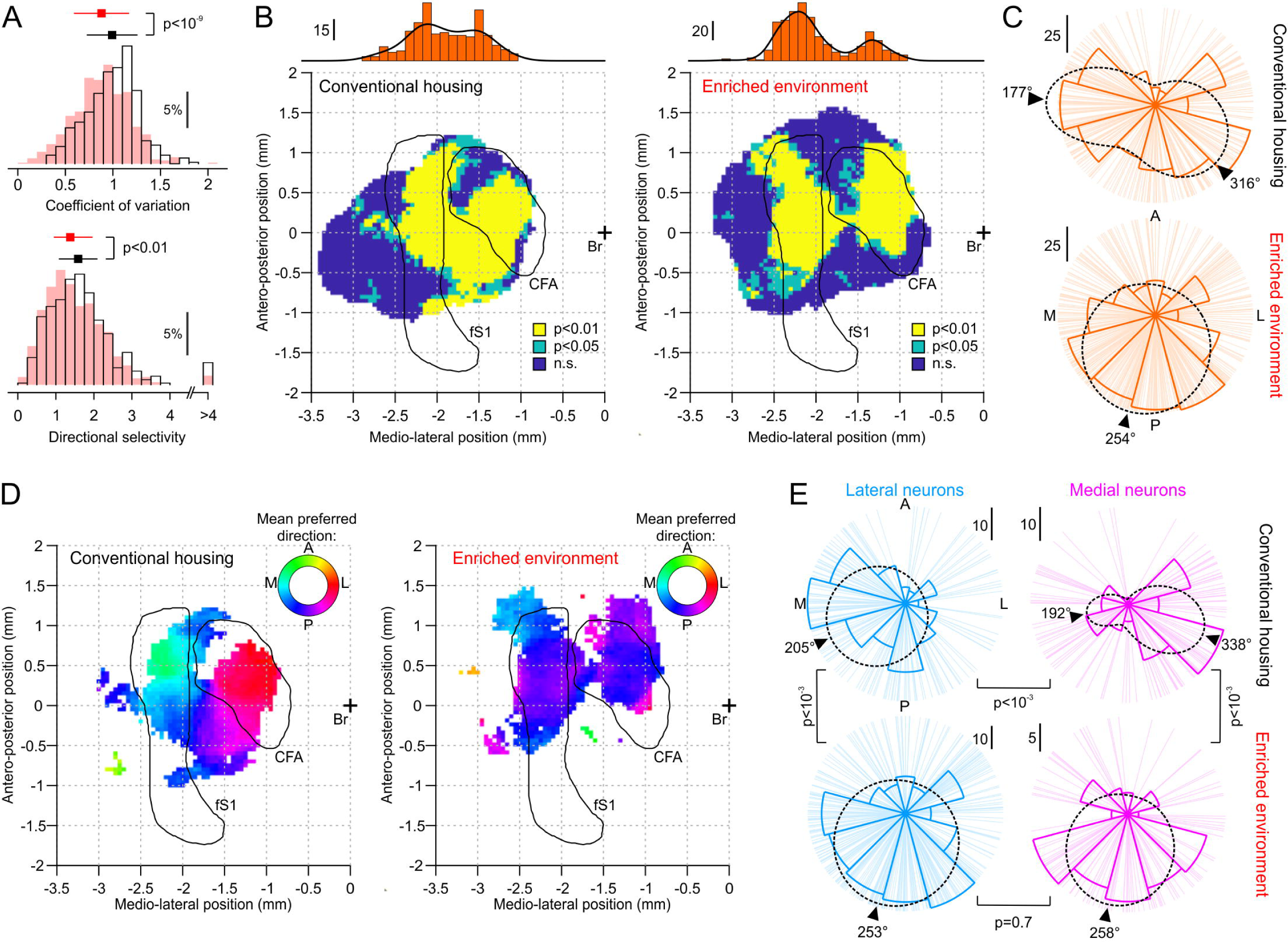
Effects of environmental enrichment on proprioceptive cortical coding. **(A)** Comparison of proprioceptive response variability and directional tuning selectivity between EE (red) and CH (black) mice. Top: coefficient of variation of peak Δf/f_0_ responses (mean +/- s.d., two-sample t-test). Bottom: directional selectivity (width of the tuning curves) quantified by the Von Mises concentration parameter κ (median +/- quartiles, Wilcoxon rank sum test). Histograms represent the distribution of these metrics for all directionally tuned neurons (EE: N=606, 5 mice; CH: N=461, 4 mice). **(B)** Topographic mapping of directional anisotropy for CH and EE mouse groups (as in Fig. 3F). Top: Distribution of medio-lateral positions for neurons located within regions of non-uniform directional tuning. (EE: N=606, 5 mice; CH: N=461, 4 mice). **(C)** Preferred direction distributions of all neurons located within anisotropic regions (non-uniform directional tuning; yellow/green areas in B) for CH (N=288, 4 mice) and EE (N=410, 5 mice) mice with von Mises distribution fits (dotted black lines: 1 distribution for EE and a mixture of 2 distributions for CH). Markers denote means of identified von Mises components. **(D)** Topographic mapping of mean preferred direction of neurons located within anisotropic regions (non-uniform directional tuning; yellow/green areas in B) for CH (N=288, 4 mice) and EE (N=410, 5 mice) groups. **(E)** Comparison of preferred direction distributions of lateral (fS1-ligned; CH: N=166, 4 mice; EE: N=303, 5 mice) and medial (CFA-aligned; CH: N=122, 4 mice; EE: N=107, 5 mice) groups (relative to the -1.75 mm lateral coordinate) of neurons in C-D (dotted black lines and markers as in C; p-values: bootstrap test on the means of dominant von Mises components).

Recomputing the anisotropy maps for the two groups independently, showed that neurons within the cortical zones exhibiting significant anisotropy displayed a pronounced bimodal distribution along the mediolateral axis in the EE group (Fig. 4B, top; p=0.001, Hartigan’s dip test), contrasting with a weaker, more overlapping one in CH controls (p=0.043). The ratio of cells within these anisotropic zones that were also located within fS1/CFA boundaries was significantly higher in EE (352/410 neurons) than in CH (182/288 neurons) mice (p<10^-11^, χ^2^test). This spatial difference suggests that, within a mesoscale activation map of unchanged size and location (Fig. 2B-E), EE aligns the cortical clusters representing proprioceptive anisotropy more precisely with anatomical fS1 and CFA boundaries (Fig. 4B). Moreover, the structure of the preferred direction distributions differed (p=0.006, Kuiper two-sample test; Fig. 4C). They were both significantly non-uniform (CH: p<10^-5^, EE: p<10^-12^, Hodges-Ajne test) and neither was strictly multimodal (CH: p=0.29, EE: p=0.26, excess-mass test). We therefore quantified the structural difference by fitting a mixture of von Mises distributions to identify the number and location of components within the anisotropic distribution (see Methods). The CH distribution was best described by a mixture of two von Mises components (ΔBIC=14.5 relative to one component; BIC: Bayesian information criterion), with components at 316° (67% of neurons) and 177° (33% of neurons) (Fig. 4C). The EE distribution was best described by a single component at 254° (ΔBIC=−4.7 for two components). A third component was not supported in any distribution (ΔBIC ≥ 1.0 relative to the selected model). These results thus suggest that EE reshapes the proprioceptive anisotropy. It seems to unify two divergent components present in CH mice into a single component at an intermediate direction, in line with the trend towards a more concentrated distribution of population activity around the mean observed with wide-field imaging (Fig. 2H).

We next asked whether CH neurons assigned to the two components were intermingled on the cortical map or spatially segregated, forming a distinct topographic organization. To test this, we first used permutation-based measures of spatial segregation (see Methods). On average, 63% of each neuron’s nearest neighbors belonged to the same component, compared with 53% expected under random intermingling (p<10^-4^, permutation test), and the centroids of the two components were 0.29 mm apart (vs. 0.05 mm by chance, p<10^-4^, permutation test), suggesting the presence of differently tuned clusters of CH neurons that map onto spatially distinct populations on the cortex. To visualize this topography, we computed the mean preferred direction across the anisotropic locations and observed a striking division along the medio-lateral cortical dimension (Fig. 4D, left). Specifically, a lateral (fS1-aligned) cluster of neurons preferred limb movements along the medio-posterior axis, whereas a more medial (CFA-aligned) cluster preferred displacements along a more lateral-posterior axis. We divided the two subpopulations by a medio-lateral cutoff at -1.75 mm relative to bregma and compared their directional preference distributions (Fig. 4E, top). The fS1-aligned cluster was described by a single component at 205° (ΔBIC=−4.7 for two components), whereas the CFA-aligned cluster had two components (ΔBIC=8.3 relative to one component), a dominant one at 338° (71% of neurons) and a minor one at 192° (29% of neurons). The dominant components of the two clusters differed significantly (p<10^-3^, bootstrap test), confirming that the fS1- and CFA-aligned clusters are tuned to different movement directions. In EE mice, this inter-areal directional divergence was absent (Fig. 4D, left; Fig. 4E, bottom). Both clusters were described by a single component (ΔBIC=−0.5 and ΔBIC=−11.2 for two components) at a common intermediate direction (fS1-aligned: 253° vs. CFA-aligned: 258°, p = 0.7, bootstrap test; reduction in the divergence by 105°, p<10^-3^, bootstrap test). Compared with CH, the dominant component of the fS1 cluster shifted by 48° (p<10^-3^, bootstrap test) and that of the CFA one by −80° (p<10^-3^, bootstrap test) (Fig. 4E). To assess whether EE also affected the concentration of preferred directions around the mean, we compared the mean resultant length r (r=0: uniformly distributed, r=1: identical directions, see Methods). The concentration within each cluster did not differ between groups (lateral/fS1: r=0.34 in CH vs. 0.26 in EE, p=0.23; medial/CFA: r=0.33 in CH vs. 0.37 in EE, p=0.67, two-sided permutation test). Thus, EE did not strengthen the anisotropy but aligned the directional preferences of the sensory and motor-aligned clusters to a common axis.

Topographic maps could not be computed from individual mouse data as each contributed on average 119 +/-27 (mean +/- s.d.) neurons, yielding too few computable windows for uniform sampling of the fS1/CFA areas and adequate statistical power. We therefore used the pooled map of each group (EE and CH) to divide each mouse’s neurons by location: inside or outside the anisotropic zones, and in the lateral (fS1-aligned) or medial (CFA-aligned) region. We then analyzed the distribution of preferred directions of each mouse’s neurons in each region (Supplementary Fig. 1A-B). The differences between the CH and EE groups identified in the pooled data (Fig. 3F, Fig. 4C, E) were consistently observed across individual mice (Supplementary Fig. 1A-E).

In summary, these results indicate that the body-petal preference in the proprioceptive cortex is topographically organized, with fS1 and CFA processing separate coordinate axes within that peripersonal space. Developmental enrichment reshapes this neural representation within a globally stable proprioceptive cortical map. It organizes the encoding of a peripersonal forelimb axis within a more precise and functionally consistent cortical topography.

## Discussion

We found that EE did not alter the mesoscale activation map of the forelimb proprioceptive cortex. Both the size and the location of the activated area were similar in EE and CH mice (Fig. 2). This contrasts with previous studies in other sensory modalities, where enrichment expanded the primary visual cortex[23] but reduced the cortical whisker representation in somatosensory cortex[24]. The three divergent outcomes may indicate that the effects of enrichment on cortical maps are modality/region-specific. Enrichment alters the amount and structure of sensory input differently for vision, whisker touch and limb proprioception, and each system may follow its own developmental time course and rules of plasticity. Alternatively, the inconsistency suggests that the size of a cortical representation is not a reliable readout of experience-dependent plasticity. Our results support this second view as the mesoscale proprioceptive map remained stable, yet enrichment reorganized its cellular properties. The most informative change was not in where neurons responded, but in which movement directions they preferred and how these preferences were arranged across the cortical surface.

The proprioceptive cortex reflects the functional properties of limb use[6]. In laboratory mice, this manifests as a preference for movements toward the peripersonal space. This is evidenced by the biased directional tuning of proprioceptive neurons[16] and the observed effects on reaching movements following their selective ablation[28]. By rearing mice in an enriched environment that promotes heterogeneous limb use (reaching-to-grasp movements, fine manipulation, climbing and running) and exploration of an expanded space, we expected this directional tuning to become more uniform. We reasoned that the proprioceptive cortex would organize during development to reflect the increased behavioral and spatial diversity by encoding a wider, more balanced distribution of movement directions. Instead, our findings reveal that enriched sensorimotor experience does not eliminate the directional anisotropy of cortical proprioceptive neurons, but that it simply improves the anatomical and functional alignment of its cortical map (Fig. 4B). The anisotropy is therefore not a consequence of impoverished limb use or motor exploration characteristic of conventional laboratory housing, but rather an ecologically meaningful property of the mouse proprioceptive cortex. We speculate that the overrepresentation of movement trajectories directed toward the body reflects an evolutionary prioritization of behaviors critical for survival, such as feeding, grooming and defensive shielding. Consequently, the cortical code may be intrinsically biased to optimize proprioceptive feedback for this peripersonal behavioral repertoire. Enriched development does not eliminate the bias, instead it allows the nervous system to support these essential movements with a re-organized topographical representation.

We observed that preferred direction distributions in fS1 and CFA are centered on different axes, implying they process limb proprioception with misaligned representations of the peripersonal space. This divergence suggests different roles for proprioceptive neurons in the two areas, for instance, state estimation vs. feedback control[29]. fS1 seems to preferentially map limb position along a body centered axis to support state estimation, whereas CFA prefers a limb-aligned axis to best use the proprioceptive feedback for movement control. The misalignment implies that communication between the two areas requires a degree of coordinate transformation. We suspect that the limited behavioral repertoire under conventional housing conditions does not require efficient sensorimotor mapping, allowing the system to tolerate the computational cost of this transformation. Conversely, the increased behavioral demands in EE mice drive the representations of both regions towards a common intermediate axis (Fig. 4D-E). By aligning the proprioceptive signals, the cortex reduces the need for additional transformations between fS1 and CFA, creating a more direct and efficient link between sensation and action. Notably, this reorganization occurred with individual neurons responding more reliably and becoming more broadly tuned (Fig. 4A). The decreased directional selectivity implies a shift toward encoding multidimensional variables rather than single features, given the theoretical principle that mixed-feature representations benefit from wider tuning curves[30, 31]. Together with elevated response reliability, the broader tuning would thus reflect optimized proprioceptive encoding of a more complex, enriched sensorimotor experience.

In conclusion, developmental experience acts primarily as a calibration mechanism for cortical proprioceptive signals. It does not overtly modify the underlying neural code. Instead, it adjusts the topography of this code to refine an intrinsic, hard-wired model of limb proprioception in the mouse cortex.

## Materials and Methods

### Animals

All experiments were performed with adult double transgenic Rasgrf2-Cre x Ai148 male mice (Jackson laboratory, #022864, Jackson laboratory #030328; 8 to 12 weeks at the start of experiments) expressing Tet-controllable GCaMP6f in layer 2/3 cortical neurons. Mice were housed in groups of five adults maximum per cage and kept, outside experimental hours, in an animal facility either under standard housing conditions (N=14, 4 used for phenotyping and 10 for imaging experiments) or a customized enriched environment (N=13, 5 used for phenotyping and 8 for imaging experiments) both under a 12h light/dark cycle. Mice were placed on a water restriction regime of 1 ml/day during imaging experiments. All procedures were approved by the Fribourg Cantonal Commission for Animal Experimentation and in accordance with the veterinary guidelines.

### Surgeries

Surgeries for chronic calcium imaging experiments were performed as previously described[16]. Briefly, mice were anesthetized, the scalp was exposed, and a titanium head frame was fixed to the skull. A circular craniotomy (∼ 5 mm diameter) was centered over the left forelimb somatosensory cortex (fS1) and caudal forelimb area (CFA). It was covered with a hand-cut cranial window glued to a stainless steel ring (5 mm OD, 4 mm ID), allowing chronic optical access for wide-field imaging. Following recovery, mice were injected intraperitoneally with trimethoprim (Sigma-Aldrich T7883; at 0.25mg/g of body weight per day over 3 consecutive days) to induce Cre recombinase dependent expression of GCaMP6f. Trimethoprim was dissolved in DMSO (Sigma-Aldrich #34869; stock solution, 100 mg/ml) and further diluted in saline prior to injection (62.6mg/ml final concentration). Behavioral experiments began 2 weeks after surgery.

### Environmental enrichment

Mice were born and housed for the duration of the experiments in a commercially available wire-mesh enclosure (Nolan, Beeztees; ∼60 × 40 × 45 cm). The cage floor was layered with standard bedding and included a small shelter (∼5 × 9 × 13 cm). The wire-mesh walls and ceiling facilitated spontaneous climbing, which was further enhanced by two elevated platforms accessible only via custom-built ladders. These ladders featured vertical rungs specifically designed to require coordinated digit grasping during ascent and descent. To necessitate regular use of this three-dimensional space and promote physical activity, standard food pellets were placed exclusively on the cage floor, while water was provided solely on one of the elevated platforms.

To promote fine motor manipulation, unshelled sunflower seeds were provided daily on the second elevated platform, and hanging ropes were suspended within the cage to elicit spontaneous interaction. Additionally, three custom-designed, motorized carousels were positioned just outside the enclosure within reaching distance. Controlled via a microcontroller (Arduino Uno) and stepper motor drivers (TMC2130 V2, Watterott), these carousels dispensed a shelled sunflower seed every 120 seconds, stimulating spontaneous reach-and-grasp behaviors.

For aerobic enrichment and monitoring, the enclosure was equipped with three running wheels integrated with infrared sensors (TCRT5000, Purecrea). Running metrics demonstrated robust engagement: individual mice used the wheels for 0.95 ± 0.54 h daily, covering a distance of 2.49 ± 1.54 km at an average speed of 2.62 ± 0.16 km/h (mean ± s.d.).

### Wide-field imaging

Wide-field Ca^2+^ imaging of population activity (Fig. 2) was performed through the cranial window with a custom fluorescence macroscope as previously described[16].

To image the Ca^2+^ activity of single neurons (Fig. 3), the macroscope was modified according to a published design[26]. The camera lenses in the imaging path were replaced with a low-magnification, high-NA imaging objective (Planapo, 5.0x, 0.5 NA, 19 mm WD; Leica 10447243), a tube lens (Thorlabs AC508-075-A-ML) in the emission path and a focusing lens (Thorlabs LA1680) in the excitation path. Continuous 470 nm excitation was focused to maximize the uniformity of illumination at the depth of cortical layer 2/3 across the field of view (∼ 3 x 3 mm). An iris diaphragm (Thorlabs SM2D25) was placed at the back aperture of the imaging objective to improve resolution at the periphery. With the used illumination geometry and the 0.5 NA lens the excitation beam is divergent when it enters the cortex, so illumination weakens with depth and keeps excitation low in the layers below the focal plane. The focal plane was adjusted for each imaging session to bring L2/3 somata into optimal focus, a single plane was acquired per session. We imaged neural activity by triggering frames (15 fps, 1024 by 1024 pixels, ORCA Flash 4.0 LT+, Hamamatsu) and writing data to disk with custom Matlab acquisition software and an Arduino microcontroller. Trials were defined by synchronizing the start and end of image acquisition with TTL markers from the behavior control system (Bpod, Sanworks). Motion correction, cell detection, ROI contour definition and temporal component extraction for each imaging session were performed with the CaImAn[32] and mesmerize-core Python libraries (github.com/nel-lab/mesmerize-core), which builds on constrained non-negative matrix factorization[33]. The model represents each neuron as a spatial footprint multiplied by a temporal trace, together with components that explicitly account for background/neuropil activity. Spatially overlapping sources, including overlap caused by projecting a three-dimensional volume onto the imaging plane, are demixed by this joint factorization. Such demixing is crucial for one-photon calcium imaging, where out-of-focus fluorescence produces a large, smooth background[34, 35]. All further analysis of the extracted relative fluorescence changes Δ*f/f_0_* for each neuron were done in Matlab.

### Proprioceptive stimulation task

Behavior was controlled and measured with real-time protocols running on the Bpod State Machine system (Sanworks). Proprioceptive stimulation of the mouse forelimb was performed following a previously described protocol[16]. Briefly, we delivered passive limb displacements to the forelimb of head-fixated awake mice using a custom robotic manipulandum. Mice were trained to grasp and maintain contact with the manipulandum endpoint while their right limb was displaced from a central home position to one of eight co-planar peripheral targets (5 mm travel distance, 3 cm/s trapezoidal velocity profile). A capacitive sensor (MPR121, Adafruit) was used to detect manipulandum releases and abort a trial. Mice were rewarded with a water droplet upon completion of a full trial without release. All eight movement directions (>10 correct trials/direction) in the horizontal plane were sampled in each imaging session.

### Indirect calorimetry and behavioral phenotyping

Metabolic and behavioral phenotypes were assessed using the Promethion Core Metabolic System (Sable Systems International) maintained within an environmental control cabinet with temperature at 23 ± 0.2 °C. Mice were individually housed and provided free access to water and food for a 5-day period. Mice were acclimatized for the first 48 h and only the last 3 diurnal (7h00 to 19h00) and 4 nocturnal (19h00 to 7h00) periods were used for analysis (Fig. 1B-D, Fig. 1H-J). For metabolic efficiency analysis (Fig. 1E-G), 24h sessions (the first 3 diurnal and nocturnal periods combined) were used for analysis. Energy expenditure or metabolic rate (expressed in kcal/h) was measured through indirect calorimetry. The system measures O_2_ consumption and CO_2_ production with high-resolution monitoring of gas concentrations in the inflow and outflow of air for each cage. Metabolic rate is calculated based on these measurements using the Weir equation. Food and water consumption were measured by weight changes using high-precision load cells. Mice were provided free access to a running wheel and distance and velocity of voluntary running bouts were measured with magnetic and optical sensors. Rearing was detected by interrupts of upper-level IR beams (outside wheel use, drinking, feeding and ambulatory periods) and their duration recorded. Ambulatory activity was detected as a sequence of IR beam breaks in the horizontal plane covering a significant distance at a velocity of > 1 cm/s. Grooming and scratching were registered as fine movements that are not ambulatory. These are detected when the mouse breaks IR beams repeatedly while staying in a fixed location. A confined habitat suspended at a 2.5 cm elevation was added to the cage and entries were detected with a high-precision load cell. Only stays longer than 10 s inside the habitat were recorded as valid entries. All measurements were taken at 15 min intervals and values averaged across each diurnal/nocturnal period. All mice had their body composition assessed at the beginning and at the end of the experimental period using an EchoMRI-100H body composition analyzer (EchoMRI LLC) and used to measure % of body fat/lean mass and normalize energy expenditure data.

### Data analysis

#### Metabolic efficiency

Metabolic efficiency was calculated as the slope of the linear relationship between energy expenditure and physical output (running distance; Fig. 1E–G). For individual 24h sessions, data points were extracted in 15-minute intervals during which wheel running activity occurred. Outliers within individual sessions were removed prior to the final regression fit using a Cook’s distance threshold of 4/n (n = number of observations). Final efficiency slopes deviating by more than two median absolute deviations (MAD) from the median were excluded from statistical analysis.

#### Wide-field population activity

To correct for shifts in macroscope alignment across imaging sessions, we registered the neural activity videos to a reference session for each mouse. We calculated an affine transformation matrix (*fitgeotransform2d.m* in Matlab) based on four landmark points on the cortical vasculature. The video frames were then aligned using *imwarp.m*. Aligned frames were then converted from pixel space to a Cartesian coordinate system (in mm, horizontal plane defined by the medio-lateral and antero-posterior axes) centered at bregma. This transformation involved scaling, translation, and rotation, determined for each mouse using bregma locations and marked anatomical distances and axes.

Stimulus aligned Δ*f/f_0_* mean traces (>80 trials/session) for each pixel location were normalized and corrected for hemodynamic signals as described previously[16]. To generate proprioceptive activity maps for each session, we analyzed the frames exhibiting peak activation following stimulus onset. Within these frames, we defined active pixels by applying a threshold to the distribution of pixel values. Specifically, we fitted a half-normal distribution to the left side of the data (the values below the distribution peak) to model the baseline noise. Pixels with values above the distribution peak plus two standard deviations of this fit were classified as ’activated’. Sessions were first averaged for each mouse prior to the computation of the group-level population activation maps (Fig. 2B).

To evaluate topographic differences in cortical proprioceptive representations between groups (Fig. 2C), we employed a non-parametric cluster-based permutation test. First, a mass-univariate t-map was generated by calculating a two-sample t-statistic at each pixel, representing the signal-to-noise ratio of the group difference:

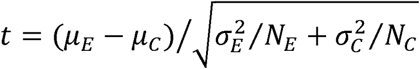

where *µ_E_* and *µ_C_* are average intensity maps of the EE and CH groups, 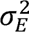 and 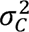 the corresponding variances and *N_E_* and *N_C_* the number of mice in each group. A threshold corresponding to p<0.05 with the given degrees of freedom (using the inverse cumulative distribution function) was applied to the observed map to identify significant difference clusters. Second, a null distribution was constructed by randomly shuffling group labels 1999 times and calculating the t-map for each permutation. We calculated the maximum cluster size and maximum cluster mass (sum of t-values) for each iteration. Cluster sizes were defined as total contiguous areas exceeding the significance threshold (expressed in mm²) and cluster masses as the integral of absolute t-values across the cluster areas (expressed in units of t.mm²), which combines spatial extent and statistical strength of the group difference. The two null distributions represent statistics expected by random chance and the p-value for each was calculated as the fraction of values greater than the maximum cluster size and mass observed for the non-shuffled data.

To compare average cortical activity along the medio-lateral and antero-posterior axes (Fig. 2E), we performed a receiver operating characteristic (ROC) analysis for every sample location along these dimensions across all sessions of the two groups (Fig. 2E). A discrimination score *D_VE_* was calculated for the *i^th^* session’s sample in the EE group as the product of its activity value (*E_i_*) and the mean value across all EE sessions excluding the *i^th^* session (E^_^) minus the product of *E_i_* and the mean values across all CH sessions (E^_^). The discrimination score *D_VC_* was obtained for the *i^th^* session’s sample in the CH group in an analogous manner:

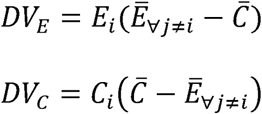

The ROC curve was constructed by plotting *p*(*D_VE_* > *k*) against *p*(*D_VC_* > *k*), the fraction of *D_VE_* and *D_VC_* values exceeding the criterion *k* varied across the range of *D_VE_* and *D_VC_* The area under the ROC curve (AUC) was computed using trapezoidal numerical integration (*trapz.m* in Matlab) and corresponds to the fraction of points correctly discriminated by an ideal observer. We repeated the analysis 999 times by randomly sampling the session with replacement and obtained a null distribution of AUC values. The differences between EE and CH groups were significant if more than 95% (p<0.05) of the null distribution values were more extreme than 0.5.

To determine the population’s preferred direction of activity (Fig. 2F-H) in each session, we computed activation maps for each of the eight stimulus directions. For each map, we identified the peak activation value and constructed a 2D polar plot, placing each peak at its corresponding movement directional angle. The preferred direction was determined by calculating the centroid of these points and identifying the vector from the origin to that coordinate.

#### Single-cell activity analysis

##### Location mapping

To map the location of imaged neurons to the Cartesian coordinate system (Fig. 3D), we applied the same registration procedure used for the population activity maps. We calculated an affine transformation by aligning vascular landmarks in the high-magnification field of view with corresponding structures in the wide-field image of the cortical window.

##### Single-neuron tuning

Proprioceptive neurons were identified by their responsiveness to passive limb displacement. We calculated the stimulus-evoked Δ*f/f_0_* response as the difference between the peak value in the post-stimulus interval (0 to 1.5 s) and the mean baseline activity (−0.75 to 0 s). To determine statistical significance (p < 0.01), we employed a randomization test. For each neuron, we generated a null distribution by calculating the response 1999 times using randomly shifted stimulus onset times across the recording trace. An observed response was considered significant if it exceeded the 99th percentile of this null distribution. To assess directional selectivity, we fitted a Von Mises distribution (circular counterpart of the Gaussian distribution) to the neuron’s stimulus-evoked responses across the eight tested directions. We validated the significance of these fits using a permutation test (N=999), where we compared the correlation between the observed and fitted data against a null distribution generated by shuffling the direction labels. Neurons with significant fits (p<0.05) were classified as directionally selective. For these neurons, the preferred direction and directional selectivity were defined as the mean μ and the concentration parameter κ of the Von Mises distribution (Circular Statistics Toolbox[36], MATLAB).

##### Anisotropy maps

The anisotropy maps were computed for pooled data (Fig. 3F) and separately for each group (Fig. 4B). A circular window (0.8 mm diameter) was moved across the cortical surface in 0.05 mm steps. At each position containing at least 10 neurons, non-uniformity of their preferred directions was tested with the Rayleigh test (p<0.05 and p<0.01). Neurons located in locations with significant non-uniformity (p<0.05) were defined as belonging to the anisotropic zones and were used for all subsequent analyses. Bimodality of the mediolateral positions of these neurons was tested with Hartigan’s dip test.

#### Statistics

To compare phenotyping metrics between groups (Fig. 1C-J), we employed a linear mixed-effects (LME) model (*fitlme.m*, MATLAB). This approach was chosen to account for the hierarchical structure of our dataset, where multiple diurnal and nocturnal sessions were nested within individual mice. Unlike standard t-tests, LME models account for the non-independence of repeated measures by treating group identity as a fixed effect and subject identity as a random effect (random intercept). This framework effectively controls for inter-subject variability while accounting for the correlation of measurements taken from the same mouse across sessions.

For the ANCOVA analysis of metabolic differences, we first tested for interactions between group and each covariate to confirm the assumption of homogeneity of regression slopes. As all interaction terms were non-significant, we proceeded with the additive model.

For all other statistical comparisons, we evaluated normality using the Kolmogorov-Smirnov test. We applied parametric Student’s t-tests to data that followed a normal distribution, while using non-parametric Wilcoxon rank sum tests for datasets that violated this assumption.

#### Circular statistics

All circular data statistics (mean, standard deviation) and corresponding statistical comparisons were computed with the Circular Statistics Toolbox (MATLAB)[36] and custom functions. All permutation p-values were computed as (1+b) / (1+N), where b is the number of permutations with a test statistic more extreme than the one observed and N the number of permutations.

##### Modality

To test the presence of more than one mode we used a circular adaptation of the excess-mass test[37]. The test statistic measures how much better the preferred directions are described by two separate arcs of high density on the circle than by a single arc.

##### Mixture components

To identify the number and location of components, each distribution was fitted with mixtures of K=1, 2 and 3 von Mises distributions by maximum likelihood. A single von Mises component is defined as a bell-shaped peak wrapped around a circle:

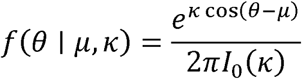

where θ is the angle (a preferred direction), *µ* the mean direction and *k* the concentration of the component, and *I*_0_(*k*) the order 0 Bessel function making the area under the curve equal one. The mixture model fitted to the preferred direction distributions adds K von Mises distributions with weights:

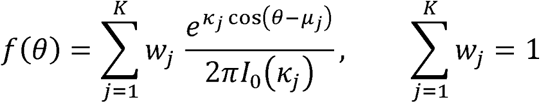

Component concentrations were bounded at *k* ≤100 and component weights at *w* ≥0.02, to prevent degenerate solutions. The number of components was selected by the Bayesian information criterion (BIC; 3K−1 parameters), and ΔBIC is reported relative to the selected model or the one-component model as indicated in the main text. The locations of the dominant components (largest weight in the selected model) were compared between clusters (lateral vs. medial) and between groups (CH vs EE, within each cluster) with a bootstrap test: p = (1+b) / (1+N). Where b is the number of bootstrap differences more extreme than the observed difference and N=1000 is the number of bootstrapped resamples of each distribution.

##### Spatial organization of components

To test whether neurons assigned to different components were spatially segregated on the cortical map, we computed for each neuron the fraction of its 10 nearest neighbors (based on Euclidean distance) that belonged to the same component. The mean of this fraction across neurons and the distance between the centroids of the two components were compared with null distributions obtained by permuting component labels across neurons (N=10000 permutations; one-sided tests).

##### Concentration

The concentration of preferred directions around their mean was quantified by the mean resultant length r. Each preferred direction was represented as a unit vector pointing in that direction, the vectors of all neurons were averaged, and r was the magnitude of the resulting mean vector. Lower values indicate a less concentrated distribution (r=1: all neurons prefer the same direction; r=0: preferred directions are uniformly distributed around the circle). To test whether this concentration differed between CH and EE mice, we compared the difference in r between groups within each cluster (lateral and medial) to a null distribution of differences using a permutation test (N=10000) with a random reassignment of group labels (CH or EE) and keeping the original group sizes.

All other statistical tests used are indicated in the main text and figure legends.

---

**Supplementary Figure 1. Directional tuning analysis on neuronal data from individual mice.**

**(A)** Distribution of preferred directions of neurons imaged in individual EE mice (columns). Neurons are categorized based on their location with respect to the topographic map of directional anisotropy computed from pooled mouse data (Fig. 4B): outside the anisotropic zones (dark blue; p-values: Rayleigh test), within the anisotropic zone (orange; p-values: Rayleigh test), in the lateral (fS1-aligned) cluster (light blue) or medial (CFA-aligned) cluster (magenta; p-values: Watson-Williams test for equal means). The fraction of directionally tuned cells (significant von Mises fit) is indicated for each mouse at the top. The fraction of cells within the anisotropic zone that are also within the anatomical boundaries of fS1/CFA is indicated for each mouse in orange.

**(B)** Same data as in (A) for individual CH mice.

**(C)** Comparison between CH and EE groups of the concentration around the mean for preferred directions of neurons within anisotropic zones (orange in (A) and (B)). Data points are individual mice (p-value: permutation test).

**(D)** Comparison between CH and EE groups of the fraction of neurons within anisotropic zones (orange in (A) and (B)) that are also within fS1/CFA boundaries. Data points are individual mice (p-value: permutation test).

**(E)** Comparison between/within CH and EE groups of mean preferred directions between/within neurons in later (fS1-aligned, light blue) and medial (CFA-aligned, magenta). Open symbols denote data from individual mice with significant lateral vs. medial differences (Watson-Williams test in (A) and (B)). P-values correspond to Student’s t-test for paired data and permutation test for non-paired data.

## Supporting Information files

**Video 1. Environmental enrichment setup.**

Mice housed in the enriched environment engaged in activities that promote forelimb use, including manipulating seeds, reaching for and grasping food, and handling ropes.

